# Boosting Single-Cell RNA Sequencing Analysis with Simple Neural Attention

**DOI:** 10.1101/2023.05.29.542760

**Authors:** Oscar A. Davalos, A. Ali Heydari, Elana J. Fertig, Suzanne S. Sindi, Katrina K. Hoyer

**Affiliations:** Quantitative and Systems Biology Graduate Program, University of California, Merced, CA, USA; Department of Applied Mathematics, University of California, Merced, CA, USA; Department of Oncology, Division of Biostatistics and Bioinformatics, Sidney Kimmel Comprehensive Cancer Center, Johns Hopkins School of Medicine, Baltimore, MD, USA; Health Sciences Research Institute, University of California, Merced, CA, USA; Department of Molecular and Cell Biology, School of Natural Sciences, University of California, Merced, CA, USA

**Keywords:** Single Cell RNA Sequencing Analysis, Deep Learning, Interpretable Deep Learning, Neural Attention

## Abstract

A limitation of current deep learning (DL) approaches for single-cell RNA sequencing (scRNAseq) analysis is the lack of interpretability. Moreover, existing pipelines are designed and trained for specific tasks used disjointly for different stages of analysis. We present scANNA, a novel interpretable DL model for scRNAseq studies that leverages neural attention to learn gene associations. After training, the learned gene importance (interpretability) is used to perform downstream analyses (e.g., global marker selection and cell-type classification) without retraining. ScANNA’s performance is comparable to or better than state-of-the-art methods designed and trained for specific standard scRNAseq analyses even though scANNA was not trained for these tasks explicitly. ScANNA enables researchers to discover meaningful results without extensive prior knowledge or training separate task-specific models, saving time and enhancing scRNAseq analyses.

---

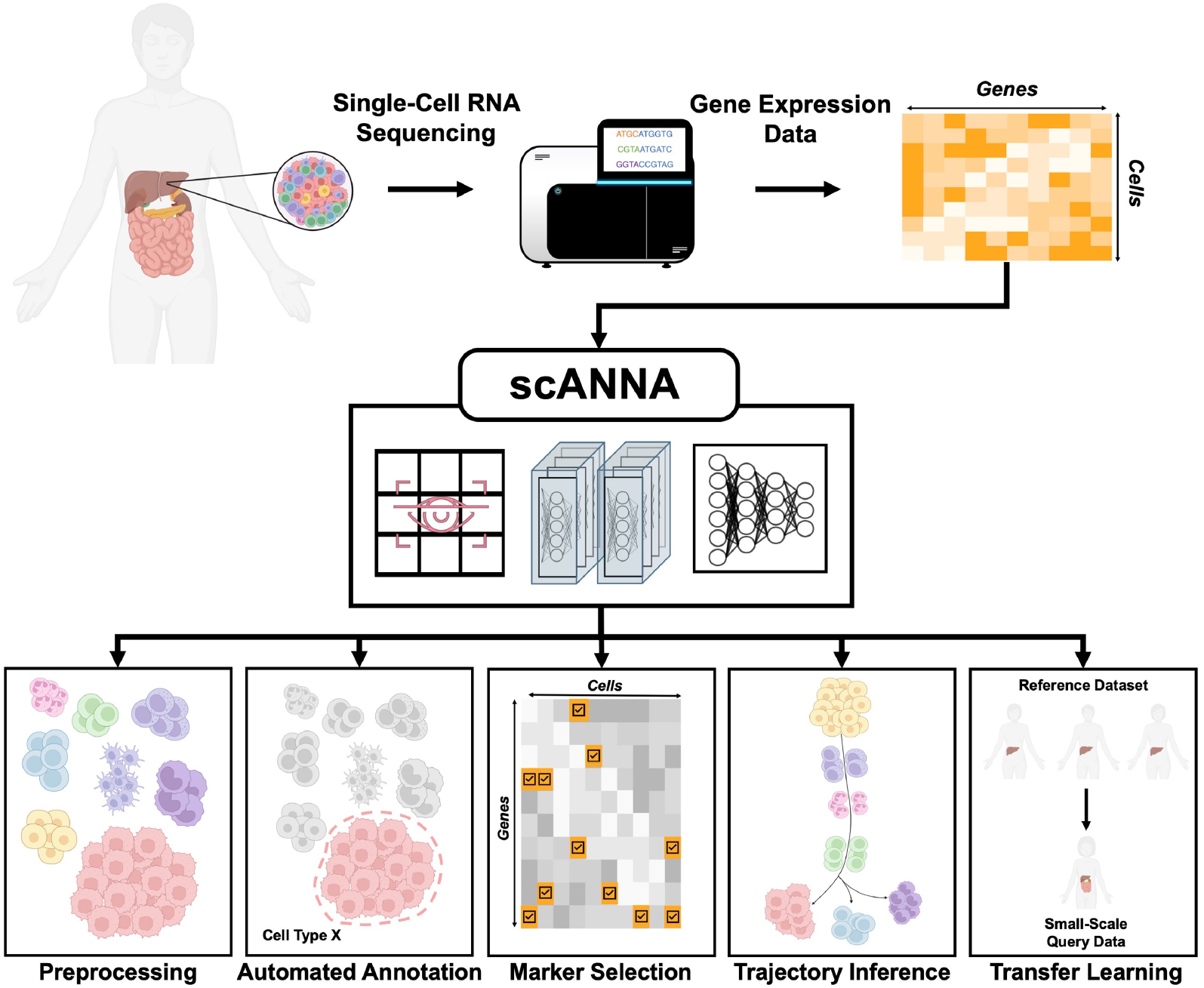

## Introduction

Single-cell RNA sequencing (scRNAseq) technologies have been instrumental in studying biological patterns and processes in development and disease (Heydari and Sindi, 2023; Szabo et al., 2019). ScRNAseq datasets are high-dimensional and noisy, often requiring advanced machine learning (ML) methods for accurate and efficient analysis (Erfanian et al., 2022). Recent years have seen a surge in deep learning (DL) methods, setting the state-of-the-art performance in many pre- and post-processing tasks, such as batch correction, dimensionality reduction, and multi-modal integration (see Erfanian et al. (2022)). While DL methods have become the standard for accuracy, a limitation of most DL approaches for scRNAseq analysis is a lack of biological interpretability; traditional DL approaches transform scRNAseq data into low dimensional representations which cannot directly be associated with specific gene signatures. Moreover, current pipelines require researchers to develop and train separate disjoint models for each specific downstream analysis pipeline. In this work, we introduce scANNA (single-cell Analysis using Neural-Attention): a flexible DL model that utilizes simple neural attention to provide unique biological insights and facilitate the discovery process through a single training procedure for a suite of downstream tasks.

The major advance of scANNA is the innovative use of a simple neural attention module within a three-module DL core (see Fig. 1**(a)**). This DL core is trained once on raw single-cell RNA sequencing counts and can subsequently be applied without retraining to various user-specified downstream tasks, such as automatic cell type classification, optimal feature selection, unsupervised scRNAseq annotation, and transfer learning (Fig. 1**(b-e)**). The DL core of scANNA has three serial modules (depicted in Fig. 1**(a)**) which take raw scRNAseq counts as input. The parameters of scANNA, i.e. the neural network weights of each module, are learned by optimizing the auxiliary objective (e.g., predicting pseudo labels) and then used without retraining on downstream tasks. For this training of the model, the auxiliary objective is designed to predict pseudo-labels, i.e. cell type labels generated through an unsupervised algorithm.

**Figure 1.**
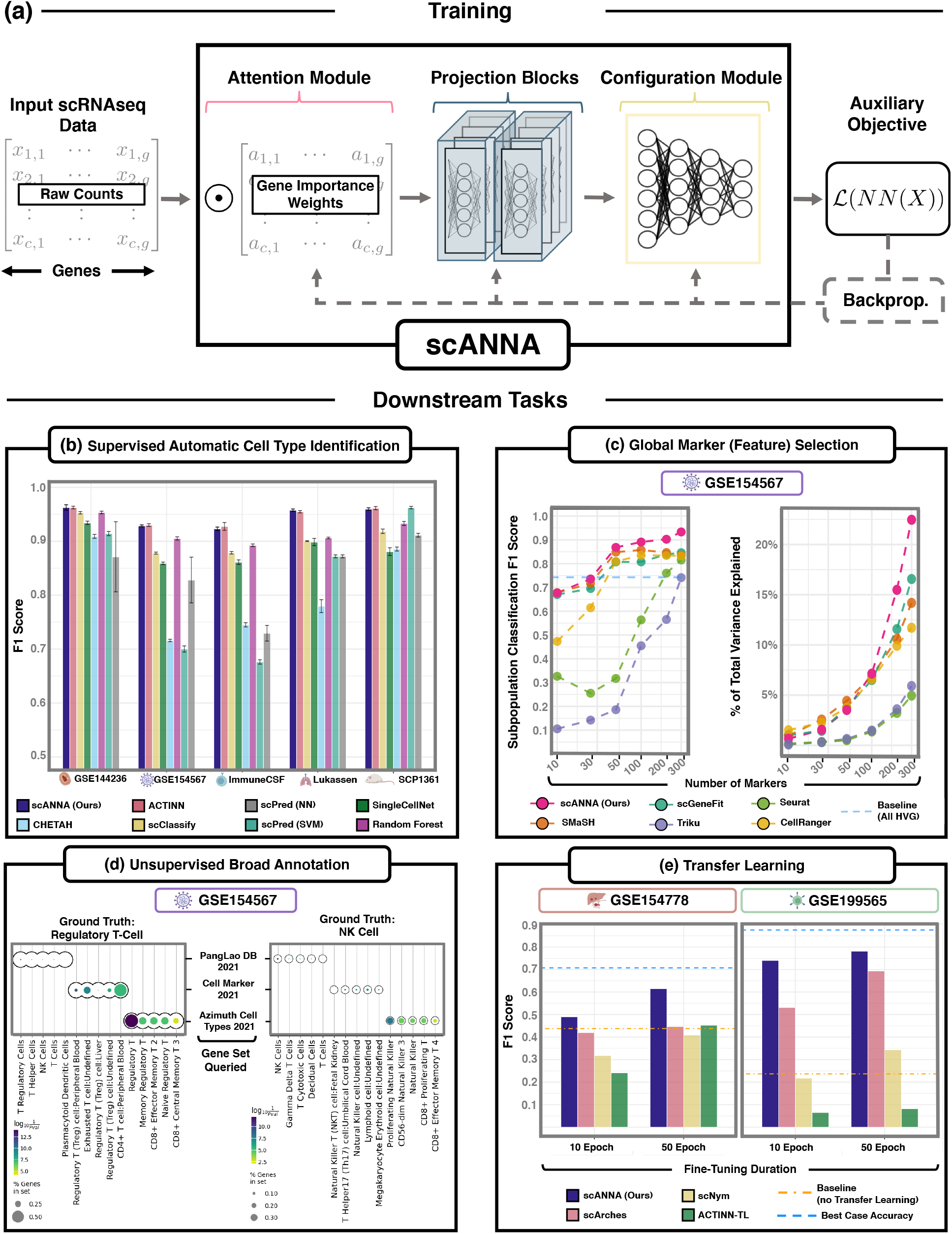
Overview of scANNA and its application to various downstream tasks. **(a)** General workflow utilizing scANNA for scRNAseq studies and illustration of scANNA’s novel three-module DL architecture. ScANNA can be used for a wide range of downstream tasks. In this manuscript, we focus on four important tasks: **(b)** Automated Cell Type Identification **(c)** Global Marker Selection, **(d)** Unsupervised Broad Annotation, and **(e)** Transfer Learning. These results show that scANNA learns complex relations from data that can be exploited for scalable, accurate, and interpretable analysis in scRNAseq studies.

The first component of scANNA is the *Additive Attention* module, which learns optimal weights for each gene based on their contribution to the auxiliary objective. These weights are used to calculate gene scores, a scaled version of the raw counts. After training scANNA’s DL core, the gene attention weights from the Additive Attention Module are used as input for most standard downstream tasks. This common training is achieved by inputting the gene scores into the second component, the Deep Projection Blocks, which are an ensemble of operators learning a nonlinear mapping between gene scores. This mapping is designed to increase model capacity and connect the gene associations to the auxiliary objective. A concatenated representation of the ensemble defined in this step forms the input to scANNA’s third component, the *Configuration Module*. The Configuration Module is the last layer of the DL core made to align with the chosen auxiliary objective function, allowing for flexibility and potential adjustments based on the dataset, (e.g. number of neurons that correspond to the number of known populations).

These weights are used to calculate gene scores, a scaled version of the raw counts. These gene scores serve as inputs to the second component, the Deep Projection Blocks, which are an ensemble of operators learning a nonlinear mapping between gene scores to scANNA’s third component, the Configuration Module. The Configuration Module is made to align with the chosen auxiliary objective function, allowing for flexibility and potential adjustments based on the dataset (e.g. number of known populations) or desired downstream applications (Online Methods). After training scANNA’s DL core, the gene attention weights from the Additive Attention Module are used as input for most downstream tasks.

In this work, we compare scANNA to state-of-the-art methods on a variety of standard scRNAseq tasks using nine public single-cell datasets. We show that scANNA attains comparable to better performance despite not being explicitly trained for these standard tasks. Based on our findings, we propose scANNA as an accurate and effective method for a variety of scRNAseq analysis settings. In large-scale studies with limited prior knowledge, scANNA’s unbiased pipelines can rapidly and accurately identify new and noteworthy genes. For small-scale studies (min 2000 cells), scANNA’s DL core can be pre-trained on existing atlases to transfer knowledge from existing repositories. As such, scANNA is a valuable tool for novice and expert users in the initial steps of scRNAseq analysis.

## Results

### Supervised Automated Cell Type Identification

While we primarily intend scANNA to be a tool for unsupervised analysis, as an initial test, we considered scANNA in a supervised learning framework. We compared scANNA against standard state-of-the-art supervised automatic cell type identification (ACTI) models ^1^ on five complex datasets (four human datasets with distinct disease conditions and one mouse dataset). ScANNA was trained on quality-controlled raw data (Online Methods), and for computational efficiency, the gene space was reduced to the 5000 (5K) most variable genes, although larger gene spaces (8K, 10K, 15K) achieve similar results (a performance drop of approximately 4% from 15K to 5K). Figure 1**(b)** and Extended Figure S3 show that scANNA performs comparably to current state-of-the-art models for ACTI (ACTINN (Ma and Pellegrini, 2019), scPred (Alquicira-Hernandez et al., 2019), SingleCellNet (Tan and Cahan, 2019), scClassify (Lin et al., 2020), CHETAH (de Kanter et al., 2019)) while providing decision making interpretability (i.e., attention tensor).

Next, we evaluated scANNA’s capabilities in two different scRNASeq tasks: global feature selection (5 datasets), unsupervised annotations (5 datasets), and transfer learning from large-scale data to small datasets (2 pairs of datasets). For each scRNASeq data set we considered, we trained scANNA once and then applied it to each corresponding downstream task without retraining the core DL model. To train scANNA in these unsupervised analysis tasks, for each data set we: (1) Generated pseudo-labels of cell types using an unsupervised clustering^2^ (2) trained the DL core to predict pseudo-labels, and (3) extracted attention weights from the Attention Module or fine-tuned the Configuration Module if transferring the learnings to a different biological context (see Online Methods).

### Global Marker Selection

We considered scANNA for global feature selection That is, we used scANNA’s learned gene weights to identify global markers, i.e. a unique and small subset of genes that are most informative and useful for additional analyses (Ma and Pellegrini, 2019). Despite the importance of global marker selection in scRNAseq studies, computational methods are nascent and often limited in the number of markers that can be selected (Alquicira-Hernandez et al., 2019). In contrast to methods that exhaustively search for n features ranked in a “one versus all” task^3^, scANNA’s Attention Module implicitly performs feature selection by assigning weights to each gene during training. After training, we extracted attention values and ranked “top” genes accordingly (Online Methods).

We compared scANNA-selected genes against markers selected by state-of-the-art ML methods (scGene-Fit (Dumitrascu et al., 2021) and SMaSH (Nelson et al., 2022), Triku (A et al., 2022)) and statistical techniques (CellRanger and Seurat) for global feature selection. Following Nelson et al. (2022), we assessed methods by evaluating classification accuracy (Weighted F1) and the fraction of total variance explained, using *n* = {10, 25, 50, 100, 200, 300*}* (Fig. 1 **(c)**, Extended Figures S5 and S6, Online Methods). Our results show that scANNA outperforms these methods in both metrics in the biologically relevant regime (between 50 - 200 genes (Nelson et al., 2022)). As such, scANNA has the potential to be a powerful addition to biological applications such as the detection of pertinent markers (100-200 genes) required for designing padlock probes used *in situ* sequencing (Heydari and Sindi, 2023; Nelson et al., 2022).

### Unsupervised Annotation

Next, we set out to perform cell type annotation, another critical challenge in scRNAseq studies (Luecken and Theis, 2019), in an unsupervised manner. A significant limitation in most computational pipelines is the reliance on manual cell type annotation based on curated lists of marker genes, which is laborious, time-consuming, partially subjective, and requires expertise (Pasquini et al., 2021; Heydari et al., 2022). In this experiment, we considered whether scANNA’s Attention Module gene weights provide a powerful companion to manual annotation. Note that as before, we use pseudo-labels for training and use the existing annotations for validation and testing purposes after training. Once scANNA was trained, we extracted the top attentive genes in each cluster as markers to query gene sets for identifying cell type, embedded in our software package (Online Methods).

Our results (Fig. 1**(d)**, Fig. 2 **(a)**, and Extended Figure S4) show that scANNA learns salient genes for each cluster enabling accurate and scalable unsupervised annotations without prior knowledge or expertise. Since many manual annotations are performed with only a few marker genes (Fischer and Gillis, 2021), it is possible to define broad and ambiguous populations; for example, two populations of GSE154567 data (see Data Description and Data Availability) annotated as “Proliferating Lymphocytes” and “Unidentified Lymphocytes”. Using scANNA’s gene scores, we disambiguate these broad annotations without re-clustering or further complex analyses. Our enrichment analysis for “Proliferating Lymphocytes” resulted in multiple significant populations, with CD8+ proliferating and effector memory T cells as the most prominent populations (Fig. 2 **(b)**). Enrichment analysis for “Unidentified Lymphocytes” yielded primarily natural killer cells with some CD8+ Effector Memory T cells (Fig. 2 **(b)**). These results signify scANNA’s utility for unsupervised annotation, and the applicability of our framework in tandem with other annotation forms to provide interpretability and validation, and scANNA performs well even with difficult-to-assign mixed populations.

**Figure 2.**
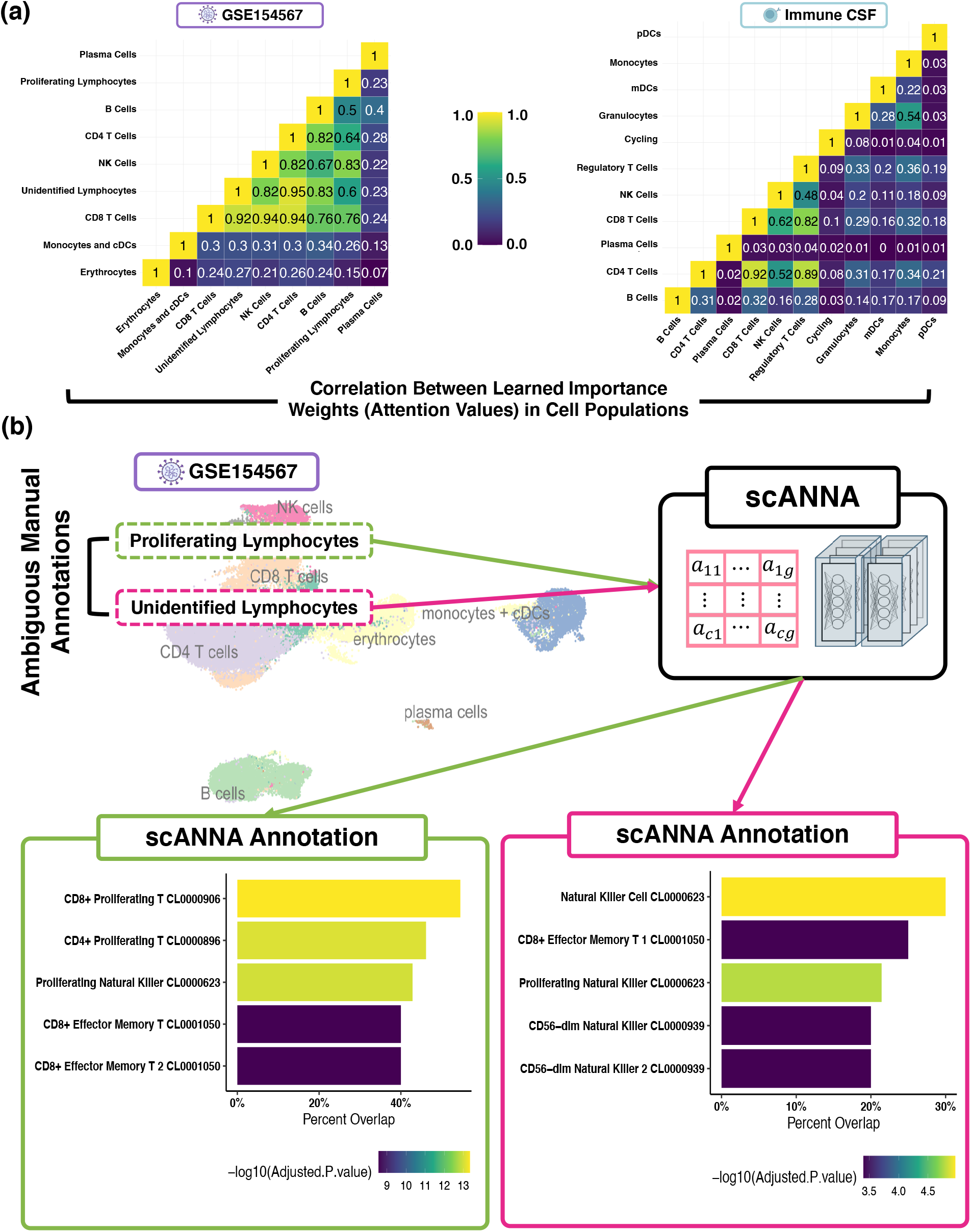
Utility of scANNA for Unsupervised Annotations. **(a)** scANNA learns complex relations between genes across cells. Here, we show correlation between attention values (learned importance weights) among different cell types in two datasets. The learned gene associations in cell types that are similar (e.g. CD8 and CD4 cells) have the highest correlations. **(b)** Enrichment analysis identified significant populations in ‘Proliferating Lymphocytes’, with CD8+ proliferating and effector memory T cells being the most prominent, while ‘Unidentified Lymphocytes’ were mainly composed of natural killer cells, along with some CD8+ effector memory T cells.

### Transfer Learning

Lastly, we considered the application of scANNA to transfer learning, the accurate and robust annotation transfer of knowledge from a larger reference set to a smaller query set. Linear machine learning methods have had broad applicability due to their interpretability (Stein-O’ Brien et al., 2019). In contrast, existing DL approaches for transfer learning (TL) on scRNSseq data typically require complex transformations prior to finetuning or fall short when the query dataset contains a different number of cell types than the source. To study scANNA’s ability to perform TL, we pretrained scANNA to predict cell types on two previously annotated atlases, (i)T cell atlas (Daniel et al., 2022) with 96750 cells and (ii) pancreatic ductal adenocarcinoma (Kinny-Köster et al., 2022) (PDAC) and 136807 cells. For target queries, we selected (i) a more specific CD8 T cell study (GSE199565) (Giles et al., 2022) with 29616 cells and (ii) a specific PDAC study (GSE154778) with 57530 (see Data Description and Data Availability).

To simulate small-scale studies and to test TL capabilities, we fine-tuned scANNA’s Configuration Module on only 20% of target datasets (5911 and 11520 cells) and used 80% for testing (23705 and 46020 cells) (Online Methods). We also analyzed scANNA performance on out-of-distribution data without fine-tuning. We compared our approach against the current state-of-the-art TL models, scArches (Lotfollahi et al., 2022) and scNym (Kimmel and Kelley, 2021). To evaluate the importance of scANNA’s architecture, we modified ACTINN for TL (ACTINN-TL; Online Methods) and used this as an additional benchmarking method. Our results (1**(e)**) show that fine-tuned scANNA achieves a higher accuracy than the other tested methods, performing within 10% F1 score of the best model (supervised classifier trained on 80% data [as opposed to 20%], denoted as “Best Case” in 1**(e)**), showing tremendous potential for TL tasks. ScANNA’s improvement over ACTINN-TL showcases the strength of attention in identifying salient genes and supporting transfer learning. ScANNA’s enhanced transfer learning has the potential to improve the accuracy and robustness of small-scale single-cell analysis.

## Discussion

We presented scANNA, a DL model employing a novel core that leverages simple neural attention for multiple scRNAseq analyses. We show that scANNA performs comparably or better than state-of-the-art methods designed and trained for specific standard scRNAseq tasks (global features selections, unsupervised annotation, and transfer learning), even though scANNA was not trained for these tasks explicitly. We have demonstrated scANNA’s effectiveness at identifying cell types in a supervised and unsupervised manner, reducing subjectivity while minimizing the time for annotating scRNAseq datasets. Our results indicate that leveraging attention in global feature selection outperforms current methods in identifying features that explain the most variance in a dataset. Lastly, we demonstrated that scANNA outperforms state-of-the-art models in TL tasks, highlighting the importance of interpretable and generalized models in the scRNAseq space. Moreover, scANNA’s unique flexibility extends beyond using cell-type pseudo-labels (as we did in this work), enabling the utilization of other groupings for learning the desired gene associations, such as leveraging hierarchies for trajectory inference.

Our results with scANNA demonstrate the potential of attention-based DL methods to offer significant improvement compared to traditional DL methods when applied to scRNAseq. Researchers can replace task-specific DL pipelines with scANNA, and potentially other interpretable DL methods that are likely to emerge. As attention-based DL methods are not yet widely used in scRNAseq analyses, our work suggests significant discoveries may yet exist in the abundance of existing public scRNAseq datasets. Finally, we note that scANNA’s architecture can be generalized to design further interpretable models for scRNAseq including spatial transcriptomics. We envision the use of attention layers in DL models will allow the field to transition from single-purpose DL models designed for specific tasks to interpretable multi-tools capable of enabling researchers without extensive computational training to accelerate the discovery process.

## Online Methods

### ScANNA: Efficient Neural Attention with Branching Projection Blocks

Our approach utilizes a simple neural attention (a feed-forward version of the additive attention introduced in Bahdanau et al. (2015)) with ensembles of linear operators, which we call *branching projections*. Multiple branching projections combined together form a *projection block*. ScANNA is a much simpler version of the popular Transformers models (Vaswani et al.) that revolutionized many fields, such as natural language processing and computer vision (Lin et al., 2021; Khan et al., 2022). Despite being simpler than Transformers, scANNA is still a large-scale model with a large capacity for learning non-linear relations in complex datasets, such as scRNAseq, while training faster than traditional transformer-based models and being easier to interpret. We describe scANNA’s different components below (visualized in Extended Figure S8):

### Additive Attention

Attention is a weighting scheme that aims to mimic the way humans understand “context” in sentences or details in images by focusing on a subset of significant features for a given objective. The use of attention-based NN for scRNAseq analysis is still in its infancy, with only a few successful scRNAseq applications to date (Yang et al., 2022; Buterez et al., 2021; Xu et al., 2021). To identify salient genes (markers), we use an additive attention module in a feed-forward NN aiming to learn the importance (weights) of all genes for each cell, given a downstream task. Analyzing the learned associations between genes and cells post-hoc provides our model’s biological interpretability.

The first step in the DL core, the Attention Module, is used to calculate a gene-score matrix (weighted version of scRNAseq count matrix), representing expression data in later layers. These importance scores enable gene prominence quantification for the downstream task, allowing interpretation of the model’s decision-making. Given a gene expression matrix *X* ∈ ℝ^*C×N*^, where *C* and *N* denote the number of cells and genes, respectively, we define the gene-score matrix Γ and the attention weights A as shown below:

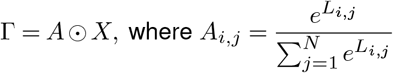

with *L* = *NN* (·) denoting a linear neural network. The learned operator *A* is leveraged after training to identify salient genes for interpretability. Gene scores have the same dimension as input data, i.e., Γ ∈ ℝ^*C×N*^).

### Branching Projections

The second module in scANNA is the projection mechanisms, which are intermediate layers between the attention layer and the Configuration Module (see Fig. 1**(a)**). The goal of using projection modules is to strike a balance between model capacity and efficiency: Too much capacity could lead to significant over-fitting, while insufficient capacity prevents the model from learning the correct representations. We design the projection blocks to allow for branching, i.e. consisting of *h ∈* ℕ separate linear operators in each level, a concept shown by Zhang et al. to improve optimization and overall learning. Such design allows efficient consideration of different gene subsets and improves model performance without the need for numerous nonlinear layers and additional computational costs. Outputs from each branch are concatenated and inputted to a subnetwork consisting of two linear operators (*L*_1_ ∈ℝ^*N ×*128^ and *L*_2 ∈_ ℝ^128*×N*^, respectively) with a Rectified Linear Unit (ReLU) in between. Through careful ablation studies, we found that *k* = 2 projection blocks with *h* = 4 branches provide the appropriate balance of accuracy and efficiency. We hypothesized that adding residual connections from the gene scores to the input of each block would improve interpretability (by reducing the chance of learning a complex unrelated non-linear mapping). Through ablation studies, we found that the residual connections, followed by LayerNorm (Ba et al., 2016) operation increased accuracy and improved interpretability. We present the architecture of a residual block in Extended Figure S8.

### Configuration Module

The last stage of scANNA consists of the Configuration Module, which is a linear operator. Our goal in separating the Configuration Module from the other components was to provide flexibility for different tasks or for transferring labels between datasets with different numbers of cells. To show scANNA’s generality, we trained one model and used a cross-entropy objective to predict labels (actual annotations in a supervised setting or pseudo-labels for the unsupervised training). The Configuration Module used for our work consisted of a linear layer mapping the gene space (number of highly variable genes) to the number of cell types, followed by a Leaky ReLU activation.

### Training scANNA

Results presented for all downstream tasks used one trained model, except for TL between reference and query datasets that had different numbers of cell populations. For all datasets, we trained scANNA by minimizing a standard cross entropy loss using the Adam gradient-based optimizer at a learning rate *lr* = 10 ^− 4^ for 50 epochs. Our experiments showed training scANNA for more epochs can result in a slight increase in prediction accuracy; however, we chose to train all models for 50 epochs for additional computational efficiency. We also employed an exponential learning rate scheduling, starting at epoch 10 and decaying every 5 epochs after with *γ* = 0.95 to avoid overfitting (more relevant when training over 100 epochs, which we did not do in this study).

### Preparation of scRNAseq dataset for analysis with scANNA

All data are publicly available from NCBI gene expression omnibus (GEO) and the Broad Institute Single Cell Portal (SCP), with links provided in Data Description and Data Availability section. Datasets were processed using the Seurat (Hao et al., 2022) package (`v4.1.0`) in R. Manual annotations were merged with count matrices using a variety of tidyverse (`v1.3.1`) functions, and subsequently added to the Seurat object as metadata. Data filtering consisted of the standard practice of removing cells with fewer than 200 expressed genes and removing genes present in fewer than 3 cells. Next, we retained cells with less than 10% mitochondrial reads to mitigate cellular debris. Lastly, cell types containing less than 100 cells were removed and excluded from the dataset. After filtering, following many similar works, we identified the top 5000 highly variable genes (HVG’s) for each dataset using Seurat’s FindVariableFeatures function, described in Stuart et al. (2019). To minimize biological and technical effects in each dataset based on patient and/or biological conditions (such as normal versus disease state), we utilized Harmony (Ko-rsunsky et al., 2019) (`v0.1.0`) to perform integration when necessary. The lower dimensional plots were generated using Uniform Manifold Approximation and Projection (UMAP) (McInnes et al., 2018). SeuratDisk (`v.0.0.0.9019`) was used to convert the Seurat data object into an AnnData object compatible with scanpy. To perform clustering and generate cell labels (in the unsupervised case), we used scanpy’s pipeline for clustering (consisting of dimensionality reduction using principal component analysis (PCA), followed by Leiden clustering). As mentioned in the main manuscript, we found Leiden resolutions that led to the same number of clusters as the annotated populations so that we could compare our predictions to the ground truth labels.

### Feature Selection

Five methods, SMaSH, scGeneFit, Triku, Seurat, and CellRanger, were compared against scANNA in evaluating its feature selection performance (i.e. selecting the most important global features (Stuart et al., 2019)). Following common metrics in this space, we measured performance by calculating the classification accuracy given top *n* features, with *n* = {10, 25, 50, 100, 200, 300}, using K-Nearest Neighbors (KNNs), XGBoost and Nearest Centroid. As a baseline, we also trained each classifier using the 5000 HVGs. In our results, we only report KNN classification accuracy, since KNN on 5000 HVGs performed best for most datasets. Additionally, we computed the fraction of variance on the original gene space (total variance) and compared that with the different approaches. Based on the evidence provided in the SmaSH manuscript and our own validation, ensemble learning (XGBoost) provided the most accurate approach, which we employed in our experiments. Of note, minor but necessary adjustments were made to the SMaSH package (located at https://gitlab.com/cvejic-group/smash) to make SMaSH compatible with `Tensorflow 2.11` (adjustments noted in the tutorial and reproducibility notebooks). For a fair comparison across all methods, we modified SMaSH to use pre-defined train and test split, which were determined a priori. Our last minor modification was to allow for the ensemble method to train in parallel (by adding `n_jobs=-1` to the `XGBoostClassifer` arguments in the `ensemble_learning` class method). For scGeneFit (Dumitrascu et al., 2021), we used the provided software package (https://github.com/solevillar/scGeneFit-python) with the same parameters as in Nelson et al. (2022) (i.e. the parameters for `get_markers` function were set to `method= ‘centers‘, redundancy=0.25, epsilon = 1`). For Triku we used the provided python package (https://github.com/alexmascension/triku) following the suggested usage protocol with only one modification (i.e. the parameter for `tk.tl.triku` function was set to `use_raw=True`). For Seu-rat and CellRanger, we leveraged scanpy’s function `scanpy.pp.highly_variable_genes` to identify important features based on the dispersion-based methods Seurat (Seurat) and CellRanger (CellRanger) (Wolf et al., 2018; Satija et al., 2015; Zheng et al., 2017).

### Supervised Classification Methods

Seven models were compared against scANNA to evaluate performance on supervised classification ACTINN (Ma and Pellegrini, 2019), scClassify (Lin et al., 2020), SingleCellNet (Tan and Cahan, 2019), CHETAH (de Kanter et al., 2019), Random Forest, scPred (Alquicira-Hernandez et al., 2019) using the svmRadial (support vector machines) model, and scPred using the neural net (NN) model. ScPred’s parameters were as follows: `resampleMethod` was set to `none, tuneLength` was set to 1, and genes containing zero counts for all cells had a pseudo count of 2 added to a cell randomly to allow scPred to run. ACTINN was run using a PyTorch implementation (https://github.com/SindiLab/ACTINN-PyTorch). Lastly, for Random Forest we used `sklearn.ensemble.RandomForestClassifier` function with default parameters. We generated five distinct train and test splits (each containing 80/20% of all data) for each dataset using the `ShuffleSplit` function from the Sci-Kit Learn package. Each model was applied to all splits with the following metric collected runtime, accuracy, weighted F1, and macro F1. Mean and standard deviation was calculated for each metric based on the dataset and model. Runtimes are presented in Extended Figures S1 and S2, and macro F1 score is shown in Extended Figure S3.

### Supervised Annotations

We chose six large immune datasets to evaluate scANNA’s capabilities across multiple diseases and tissues. Five datasets consist of human cells and one of the mouse cells. The human datasets included three SARS-CoV-2 viral infection studies and a human cutaneous squamous cell carcinoma study. We included a mouse dataset to demonstrate our model’s effectiveness on non-human and non-immune datasets. All datasets were generated using the 10X Genomics platform. A summary for each dataset is provided in Data Description and Data Availability section.

### Unsupervised Annotations

Given that labels are not available in an unsupervised setting, pseudo-labels are used to keep the same supervised objective as in the dataset publications. Pseudo-labels can be generated using traditional methods, such as graph-based clustering, or self-supervised contrastive learning (for which scANNA can also be used). To avoid any improvements in annotation not caused by scANNA, we generated pseudo-labels for each dataset using the same clustering approach described in the originating manuscript. After clustering, the arbitrary cluster numbers were used as pseudo-labels, and scANNA was trained to predict the cluster number given to a cell. No information about the cell types, known marker genes or common housekeeping genes were used during training. After training, we extracted the top n attentive genes and used those genes as markers for querying cell type. using GSEAPY’s (https://github.com/zqfang/GSEApy) enrichr method. We provide an overview of our workflow in Extended Figure S4.

### Utility of scANNA for Disambiguation of Broad Annotations

Given that many manual annotations are typically performed with only a few genes (marker genes) (Pasquini et al., 2021), it is possible to have populations that are broad and ambiguous. This was the case with two populations of GSE154567 data (Data Description and Data Availability): The original annotations “Proliferating Lymphocytes” and “Unidentified Lymphocytes” (Fig. 2 **(b)**). Our goal was to utilize scANNA to disambiguate these broad annotations without re-clustering or performing other complex analyses. Therefore, we investigated the two broad annotations in GSE154567 data and queried our top 50 attentive genes from the Azimuth Cell Type 2021 database33 using Enrichr R Package (v3.0) (Chen et al., 2013; Kuleshov et al., 2016; Xie et al., 2021) to perform enrichment analysis.

Our enrichment analysis for “proliferating lymphocytes” resulted in multiple significant populations, with the most prominent type being CD8+ proliferating T cells (2 **(b)**). Intuitively, these results are expected given the quantitative and qualitative similarity of this population with the CD8+ T cell population. The difference in gene overlap percentages of the top three predictions, namely CD8+ proliferating T, CD4+ proliferating T, and proliferating natural killer cells is very small due to the similar lineage and function of these populations, which may explain why the original annotations were left broadly as “proliferating lymphocytes“. Enrichment analysis for “unidentified lymphocytes” yielded natural killer cells as the most probable cell type, with other viable populations also being statistically significant. Similar to the previous population, these results are intuitive since CD8+ T cells and natural killer cells are both lymphocyte subsets and proliferate. There is extensive overlap in the gene sets between CD8+ T cells and natural killer cells due to their similar lineage and function. Lastly, we note that the second enriched population based on attentive genes is CD8+ effector memory T, suggesting that the “unidentified lymphocyte” population could be refined into two or more populations. These results further signify the utility of scANNA for unsupervised annotation, and the applicability of our framework in tandem with other annotation forms to provide interpretability and validation and the ability to identify functional linage differences between similar cell subsets that can be exploited in experimental studies.

### TL Experiemnts

For transfer learning, we focused on labeling two small-scale query datasets using larger source data (references). These experiments were as the following:

#### Pancreatic ductal adenocarcinoma (PDAC) TL

As an initial assessment for transfer learning, we pre-trained scANNA on a PDAC atlas, an integration of five publicly available datasets (see Data Description and Data Availability), which were split 80% and 20% for training and testing, respectively. As the target data, we chose Lin et al. (GSE154778) (Lin et al., 2020) since it contained scRNAseq of the pancreatic primary tumor while containing important distinctions and differences from the source data, making it appropriate for transfer learning. From this data, we used 20% of randomly-selected cells for fine-tuning, and 80% for testing, aiming to mimic a small-scale study. Note that the fine-tuning process uses the reverse training/testing proportions of pre-training splits to test the performance of the TL experiment.

#### T cell atlas to CD8+ T cell Populations

As a more challenging task, we aimed to perform TL from a broad T cell atlas (GSE188666) to a CD8+ T cell-specific dataset (GSE199565), which contains more subpopulations than the reference data. Similar to the PDAC experiment, we used 80% of the source data for pretraining (and 20% for testing the pre-training), 20% of the query target data for fine-tuning, and 80% for testing (see Data Description and Data Availability).

### Fine-Tuning scANNA

Fine-tuning scANNA consists of freezing all weights in the attention and projection modules, with the only trainable parameters being in the configuration module (the last layer of scANNA mapping projections to the corresponding cell types). We fine-tuned scANNA (and other tested models) for 10 or 50 epochs using the same hyperparameters as pre-training.

### scArches for TL

ScArches (Lotfollahi et al., 2022) allows transferring labels to query data through integrating it with a reference atlas. ScArches relies on an underlying reference-building method. For our application, the appropriate underlying models are scANVI (Xu et al., 2021) and scGen (Lotfollahi et al., 2019) as described in (https://scarches.readthedocs.io/); however, we were unable to use scGen due underlying maintenance issues, therefore all results presented in the manuscript are for scArches with scANVI (which uses scVI at its core). Though scGen method is recommended, we note that scANVI is also an appropriate method since it can be trained unsupervised and finetuned on a target data using labels, thus making the comparison relevant and fair. Using the source data, we trained the scVI core for 100 epochs, with an additional 10 epochs for the annotation which achieved desirable accuracy (comparable to other methods). Then, similar to scANNA, we fine-tuned the model for 10 and 50 epochs on the query data.

### ACTINN-TL

We hypothesized that scANNA’s advantage for TL stems from our architecture design. To test the importance of architecture, we applied our methodology for freezing weights and fine-tuning to an existing model, ACTINN, with the following rationale: If scANNA’s architecture is not unique and more beneficial for transfer learning, then a pre-trained ACTINN model (on the source data) should perform comparably to scANNA (once fine-tuned on a query dataset) given its comparable performance in the supervised annotation regime. Similar to the other experiments, we trained ACTINN-TL on source data for 50 epochs, then froze the weights in the hidden layers, and fine-tuned the last layer responsible for predicting the correct cell type. For simplicity, we call this approach ACTINN-TL.

### scNym for TL

ScNym (Kimmel and Kelley, 2021) model is a semi-supervised method combined with MixMatch framework (Berthelot et al., 2019) and domain adversarial training for transferring labels from a source to a target dataset. scNym generates pseudo-labels for target cells, and randomly pairs source and target observations with the weighted average of each being computed. ScNym then aims to minimize a supervised classification loss on the paired “mixed” training examples while minimizing the interpolation consistency loss on the “mixed” target cells. We trained scNym with the source and target data using the default setting for 100 epochs and then continued training for additional 10 or 50 epochs with the pre-trained model using the test target data.

### Data Description and Data Availability

#### SCP1361

SCP1361 consists of aortic cell scRNAseq from mice fed a normal diet or a high-fat diet, resulting in 24K cells (after processing) (Kan et al., 2021). The authors identified 27 clusters, for 10 different cell populations. Original data for SCP1361 can be downloaded from Single Cell Portal https://singlecell.broadinstitute.org/single_cell.

#### ImmuneCSF

ImmuneCSF (GSE163005) profiles scRNAseq data in cerebrospinal fluid (Heming et al., 2021). Cells isolated from 31 patients: 8 Neuro-COVID patients, 9 non-inflammatory, 9 autoimmune neurological diseases, and 5 viral encephalitis resulting in a total 70K cells after processing, with 15 populations. Raw data can be downloaded from Gene Expression Omnibus (GEO) with accession number GSE163005.

#### GSE154567

GSE154567 evaluates the transcriptional immune dysfunction in patients triggered during moderate and severe COVID-19 using scRNAseq (Yao et al., 2021). Peripheral blood mononuclear cells were isolated and sequenced from 20 patients. Patients ranged from healthy (*n* = 3), moderate COVID (*n* = 5), acute respiratory distress syndrome (ARDS-Severe, *n* = 6), and recovering (ARDS-Recovering, *n* = 6), resulting in 69K cells. After pre-processing the dataset, we retained 64K cells. The authors identified 9 populations in the dataset. Raw data can be downloaded from Gene Expression Omnibus (GEO) with accession number GSE154567.

#### GSE144236

GSE144236 evaluates scRNAseq of normal skin and cutaneous squamous cell carcinoma (cSCC) tumors (Ji et al., 2020). Normal and cSCC tumor cells were sequenced from each patient (*n* =10), resulting in 48K cells and 7 major cell populations. We retained 47K cells after pre-processing. The myeloid cell population (CD14+Hi) is composed of various sub-populations. Raw data can be downloaded from Gene Expression Omnibus (GEO) with accession number GSE144236.

#### Lukassen Lung

Lukassen (EGAS00001004419) evaluates single nuclei RNA sequencing of lung tissue to evaluate the expression of ACE2 and TMPRSS1 (Lukassen et al., 2020). Primary lung tissue was sampled from male and female smokers and non-smokers (*n* = 12 total), resulting in 39K cells and 9 cell populations. We retained 39K cells after pre-processing. Raw data is available from the European Genome-Phenome Archive (EGA) with study id EGAS00001004419.

#### PDAC atlas

PDAC atlas consists of five scRNAseq published datasets (Steele et al., 2020; Elyada et al., 2019; Moncada et al., 2020; Bernard et al., 2019; Peng et al., 2019) from normal and pancreatic ductal adenocarcinoma patients. After pre-processing and integration (as described by Kinny-Köster et al. (2022)), we retained 85,437K cells with 11 subpopulations.

#### PDAC small

PDAC small (GSE154778) is a specific PDAC study consisting of individual cells (57K with 10 subpopulations) from dissociated primary tumors or metastatic biopsies obtained from patients with PDAC (Lin et al., 2020). Raw data can be downloaded from Gene Expression Omnibus using accession number GSE154778.

#### T cell atlas

T-cell atlas (GSE188666) evaluates CD8 T cell exhaustion during viral infection by scRNAseq resulting in 96K cells, (retained 96K cells after preprocessing) (Daniel et al., 2022). Raw data can be downloaded from Gene Expression Omnibus (GEO) with accession number GSE188666.

#### GSE199565

CD8 specific data (GSE199565) evaluates CD8+ T cell temporal differentiation by scRNAseq resulting in 29K cells (retained 29K cells after pre-processing) (Giles et al., 2022). The authors identified 19 CD8 T cell populations (many populations were intermediate populations). Raw data can be downloaded from Gene Expression Omnibus (GEO) with accession number GSE199565.

## Code Avalibility

Our code package, including all source code and dependencies is publicly available at: https://github.com/SindiLab/scANNA. Reproducibility scripts and links to comprehensive tutorial notebooks are provided at https://github.com/odavalos/DavalosHeydari_scANNA. To improve user experience and reproducibility, we actively monitor the repository for any issues or improvements suggested by the users.

## Acknowledgements

The authors would like to thank Erica Rutter for their valuable feedback and suggestion in the early stages of this work. A.A.H is supported by the National Human Genome Research Institute of the National Institutes of Health under award number F31HG012718 (National Research Service Award). O.A.D was supported by an NIH diversity supplement R15-HL146779. The study was supported by the National Institutes of Health (R01-GM126548), National Science Foundation (NSF) (DMS-1840265) and University of California (UC) Office of the President and UC Merced COVID-19 Seed Grant. Computational resources were supported by the NSF, Grant No. ACI-2019144 and in part through NSF awards CNS-1730158, ACI-1540112, and ACI-1541349. Part of this research was conducted using Pinnacles (NSF MRI, # 2019144) at the Cyberinfrastructure and Research Technologies (CIRT) at the University of California, Merced. Our abstract figure contains graphics from BioRender, which are used under the license agreement WS257WUMFP.

## Contributions

A.A.H conceived the project with guidance from E.J.F and S.S.S. A.A.H and O.A.D implemented the algorithms, designed experiments, and performed evaluations. O.A.D, A.A.H, and K.K.H selected the biological datasets, and O.A.D performed data preprocessing. E.J.F., K.K.H, and S.S.S. provided feedback on the methodology and results. All authors helped in writing and revising the manuscript.

## Extended Figures

**Figure S1.**
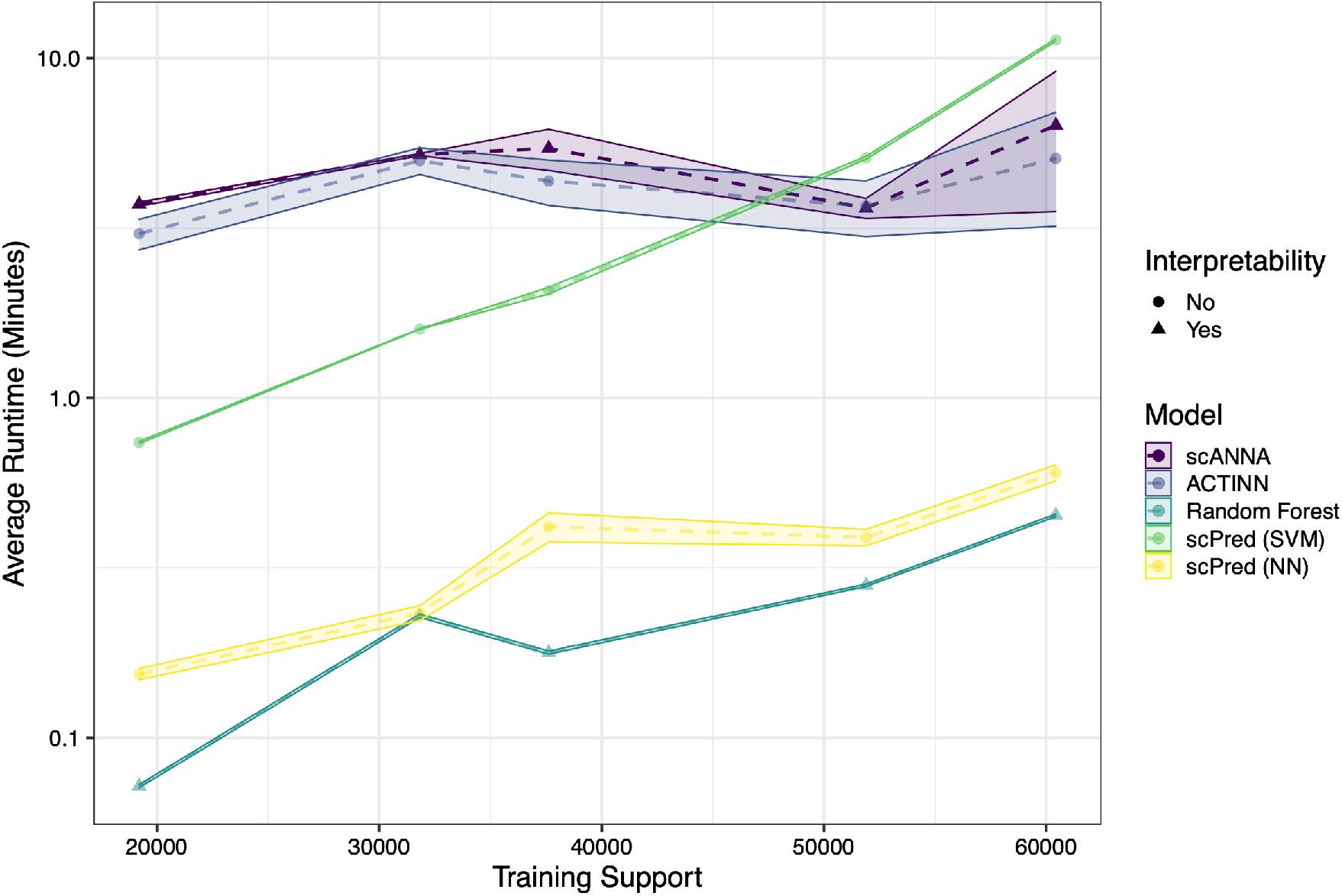
Comparison of average model training times when utilizing GPUs for deep learning models. Although scANNA is a much larger model than ACTINN (∼760M parameters compared to ACTINN’s 25M), scANNA trains in a comparable time as ACTINN. Note that Random Forest and scPred were trained on CPUs (same runtimes as Extended Figure 2) and are used here as baselines.

**Figure S2.**
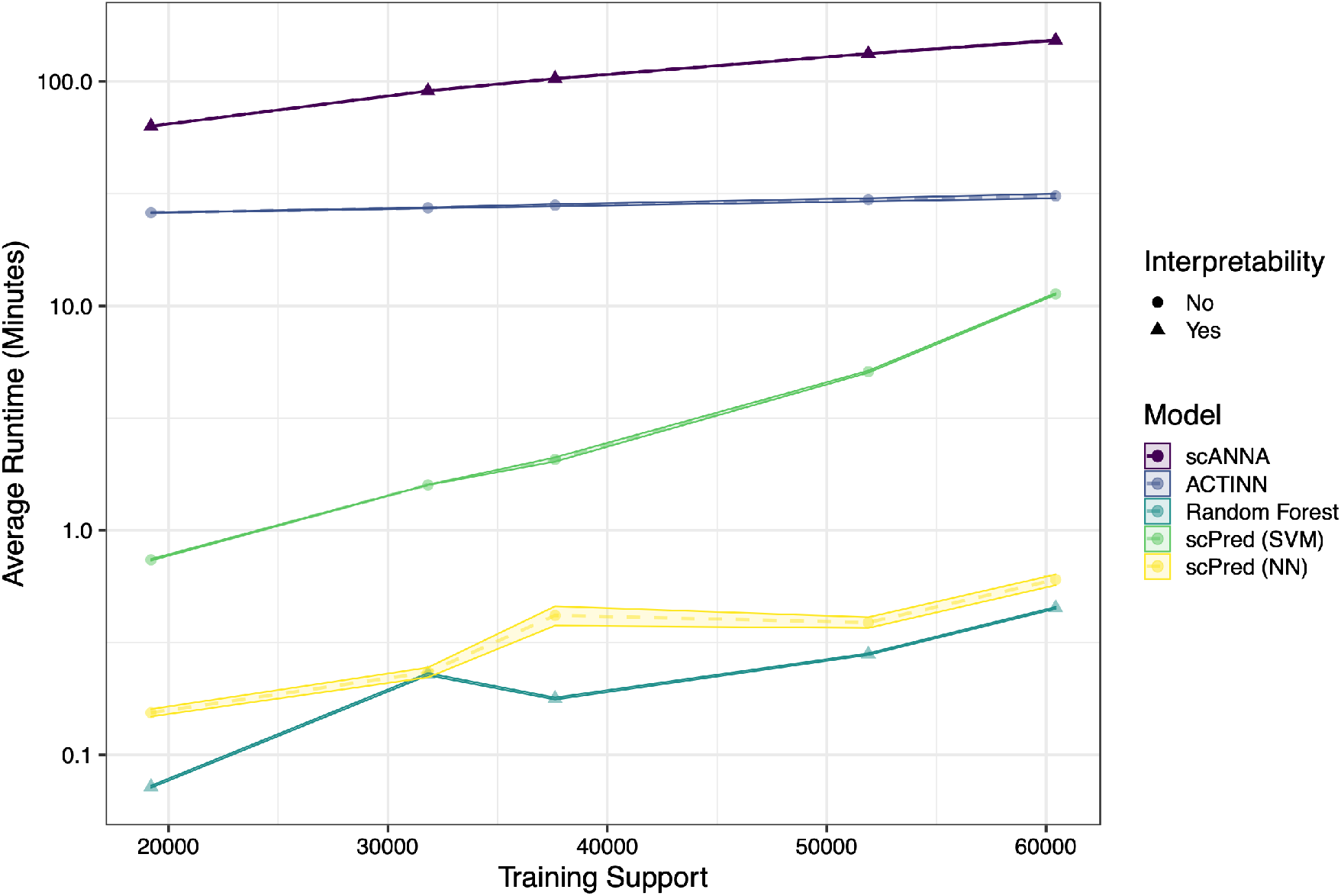
Comparison of average model training times when utilizing CPUs. ScANNA takes considerably longer time to train on a standard computer (provide specifications here) than traditional methods. However, once trained, scANNA can be quickly finetuned on other datasets for downstream analyses (10*×* reduction in runtime on average).

**Figure S3.**
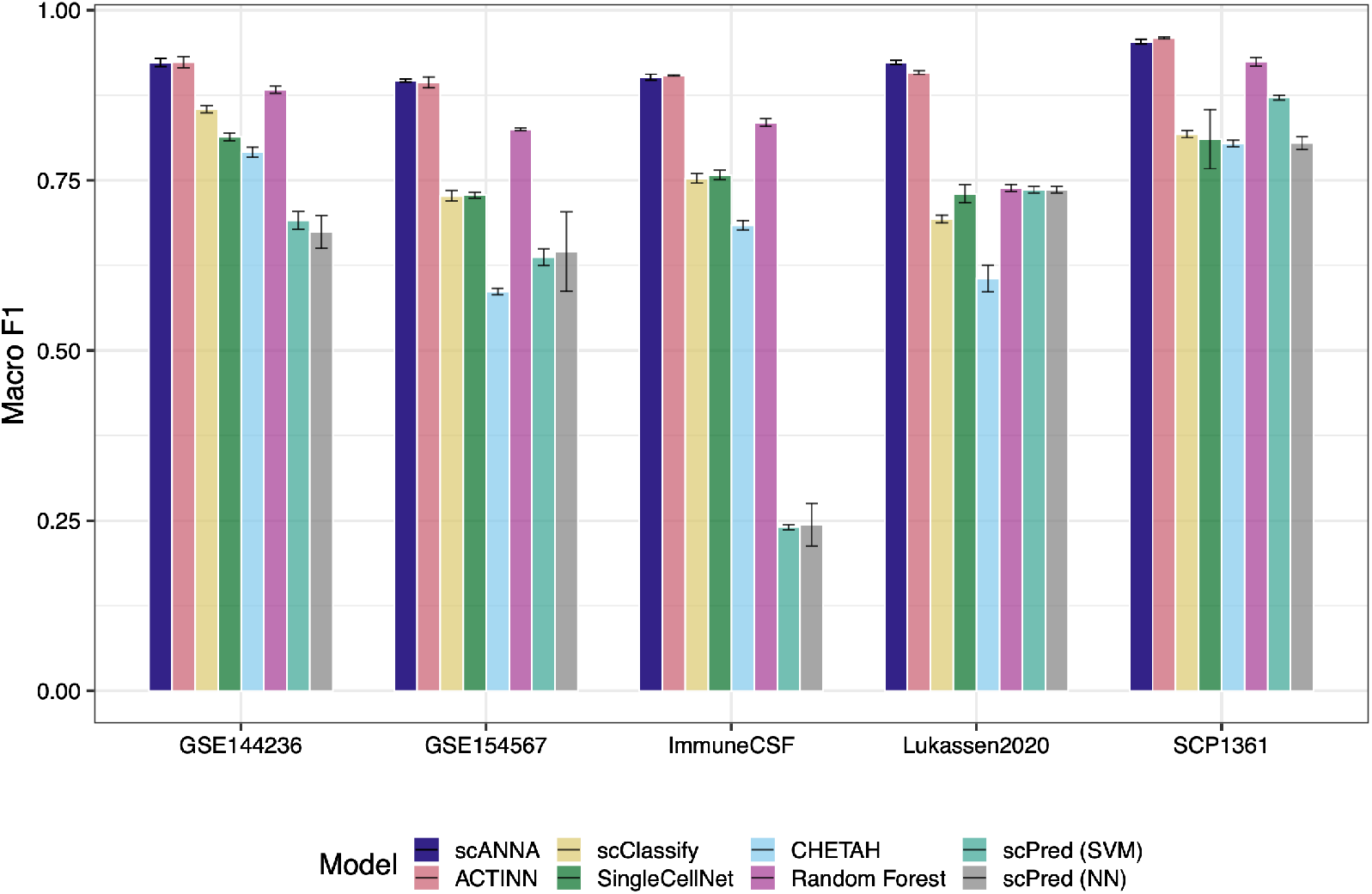
Accuracy of various supervised annotation methods reported in macro F1 score.

**Figure S4.**
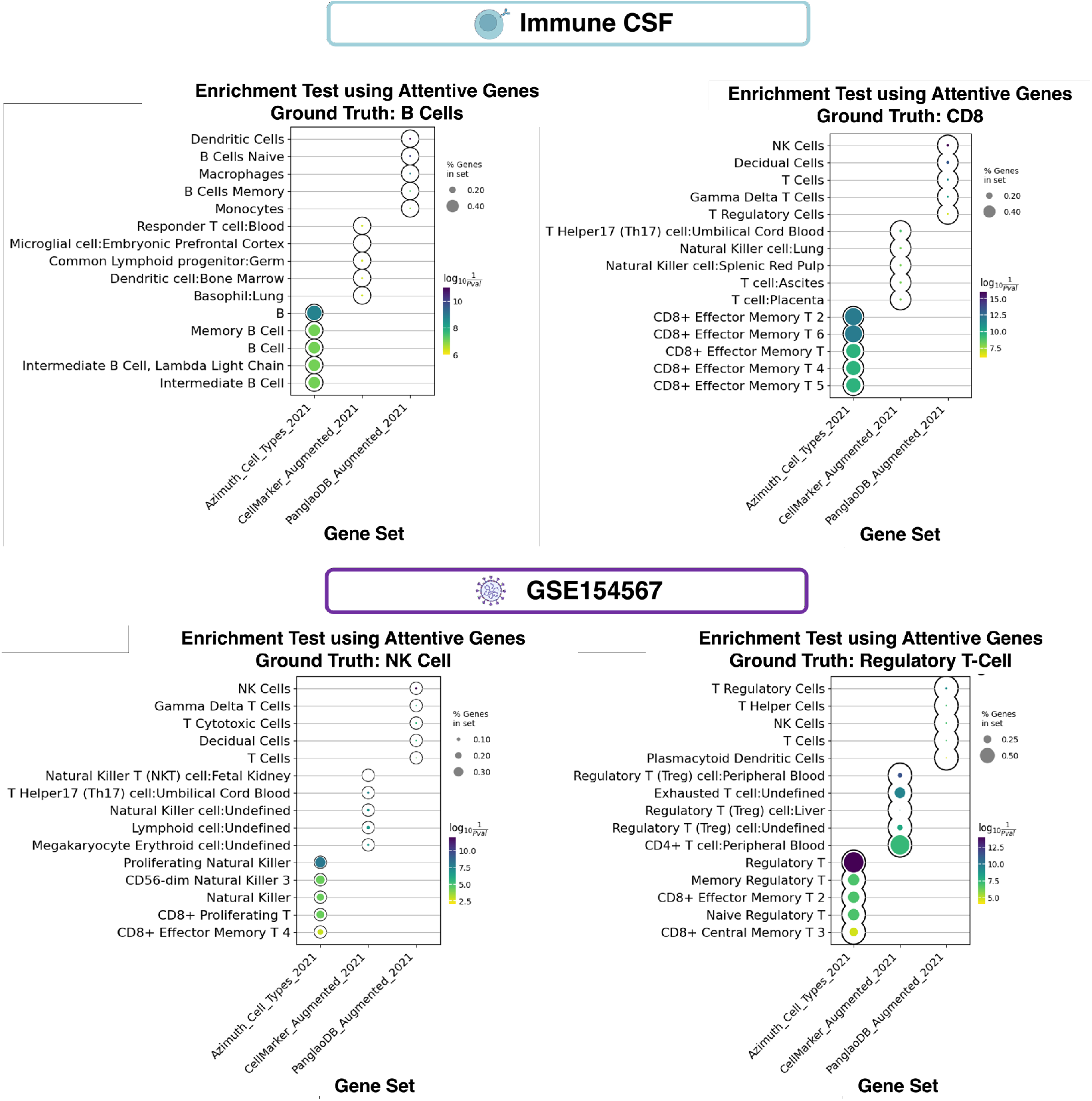
ScANNA’s unsupervised broad annotation example (enrichment of local marker genes identified by scANNA).

**Figure S5.**
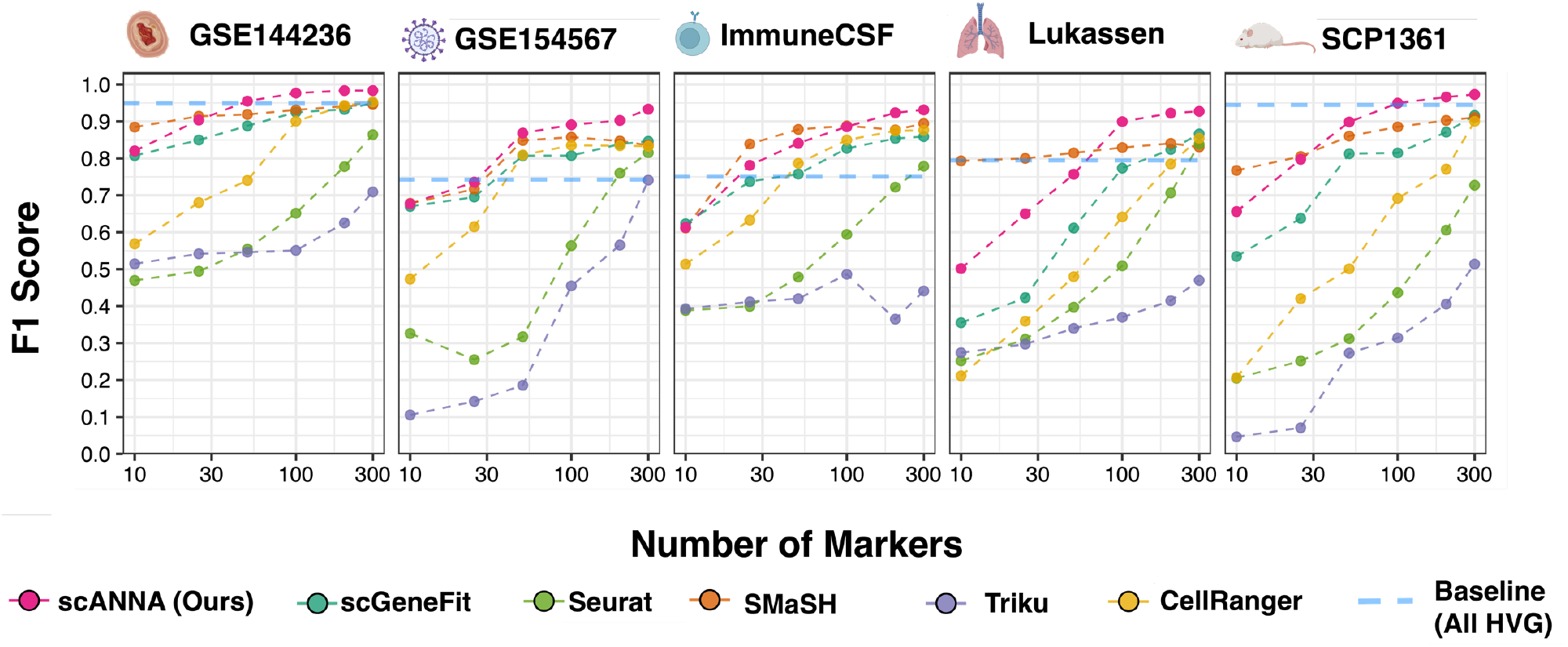
Evaluating global marker selection performance through measuring classification accuracy of cell populations with *n* selected markers from each model.

**Figure S6.**
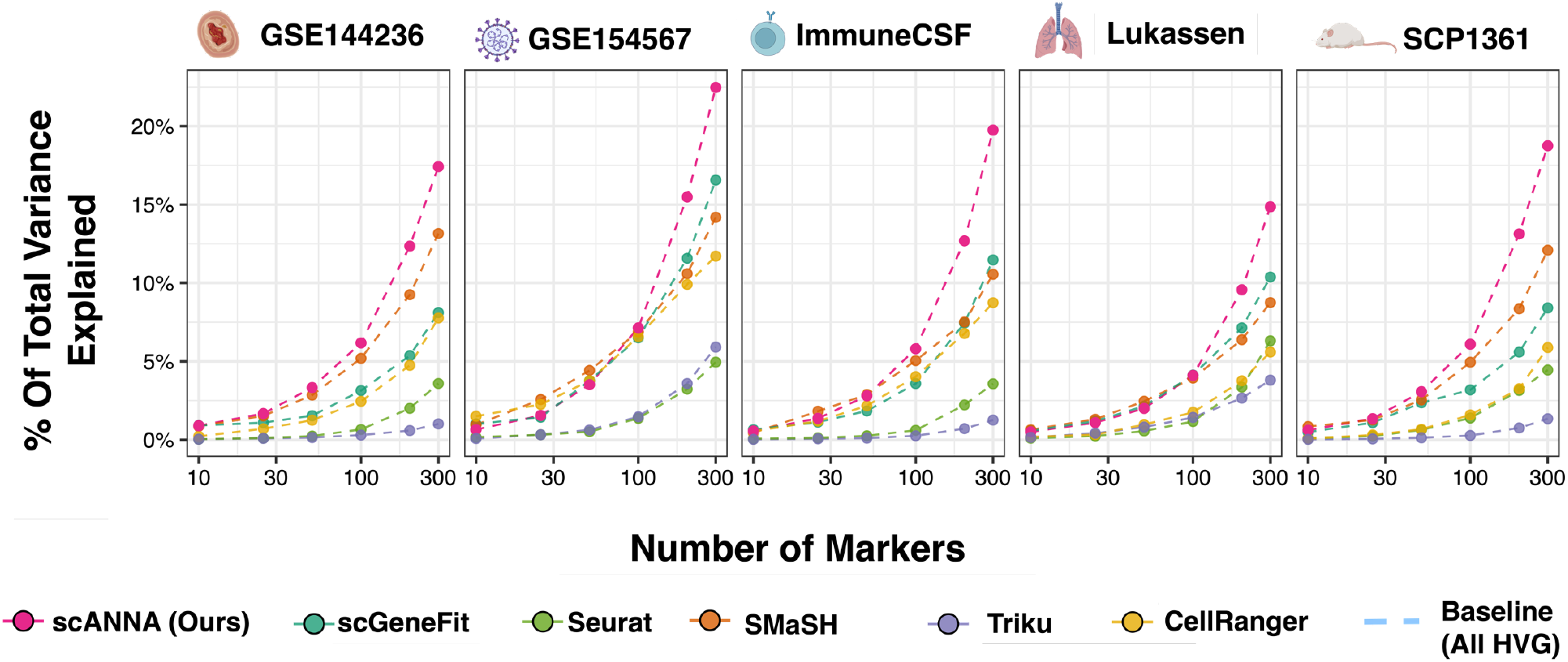
Evaluating global marker selection performance through analyzing the fraction of total variance explained with *n* selected markers from each model.

**Figure S7.**
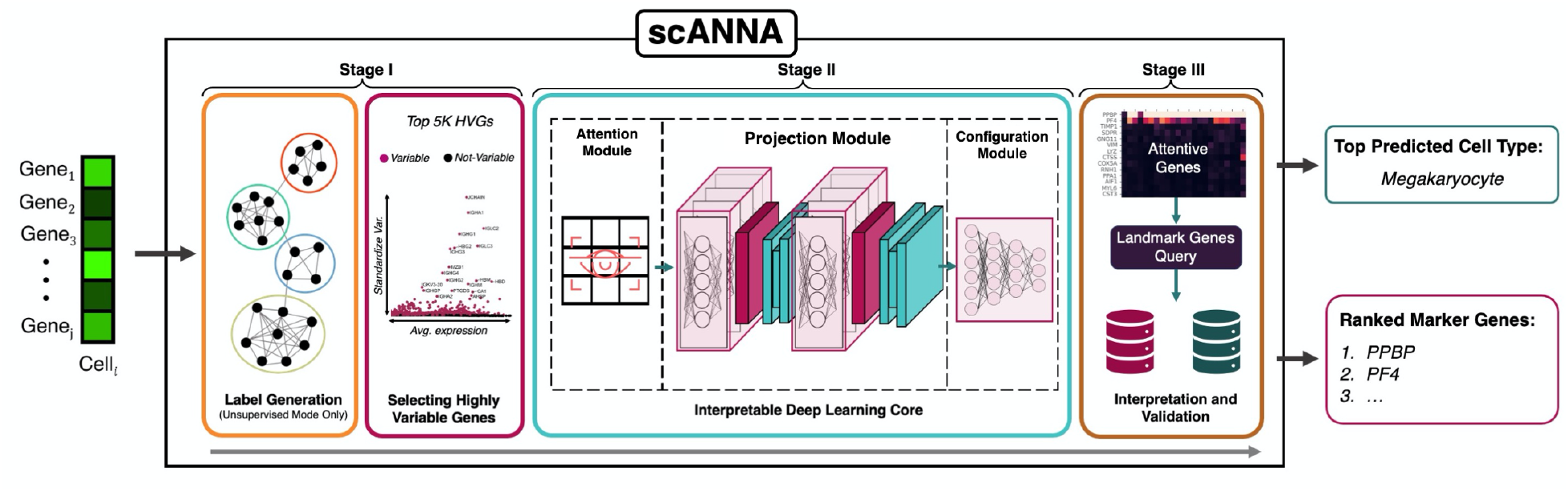
Overview of scANNA’s workflow of performing unsupervised annotation used in this study.

**Figure S8.**
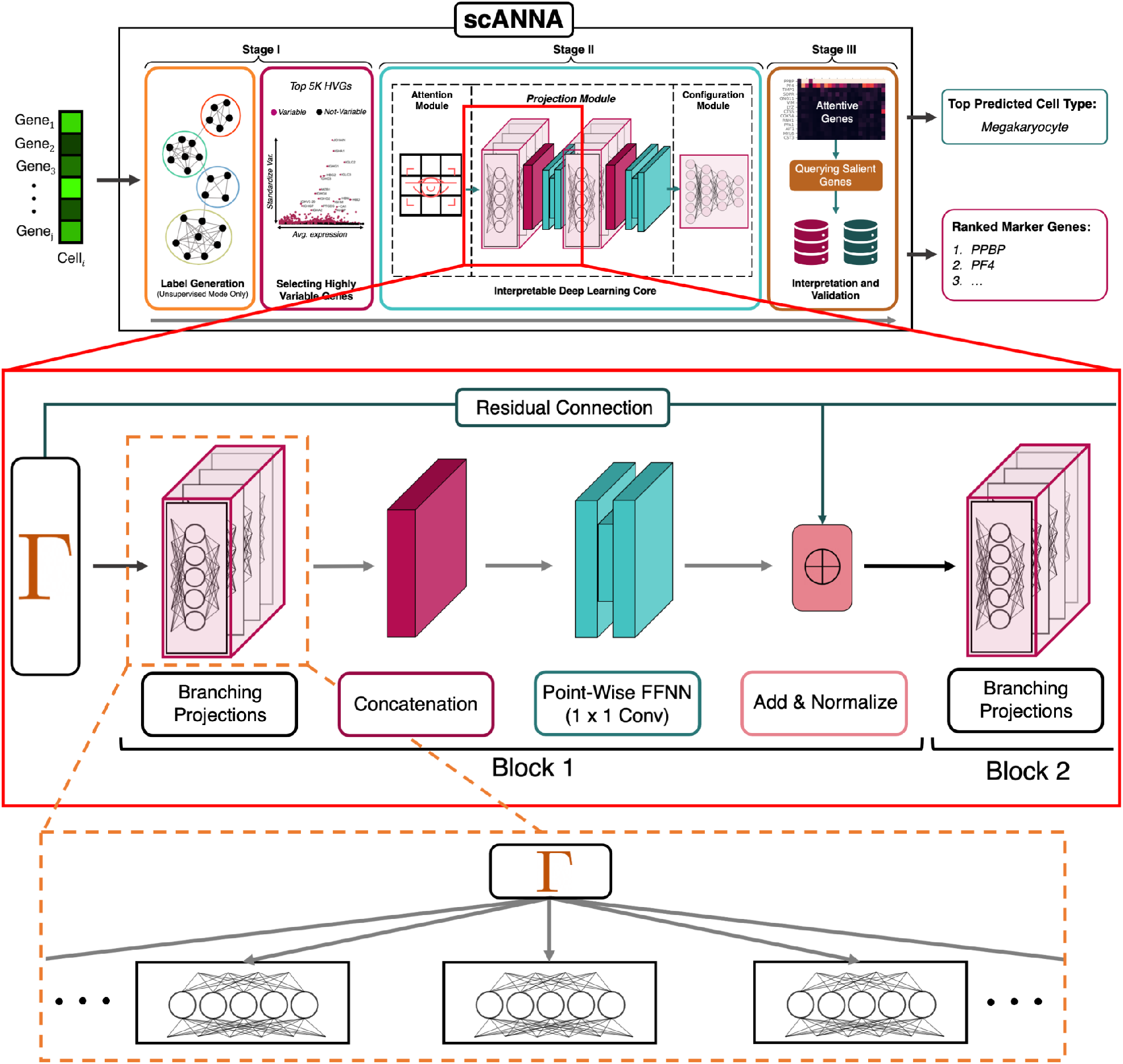
Overview of scANNA’s projection block with respect to other components (using the unsupervised annotation as an example).

## Footnotes

1 Although this is not scANNA’s primary application. Here we use the ground truth labels, in the other studies we use only our generated pseudo-labels.

2 While it is possible to use more advanced methods, such as self-supervised contrastive algorithms, we chose a naїve unsuper-vised clustering process in order to assess improvements offered by scANNA more directly. For the 5 datasets used in feature selection and unsupervised annotation, we conditioned the number of clusters to be the same as the number of manually-annotated populations for a fair comparison.

3 One v *n* marker selection refers to evaluating the importance of each gene against n other genes (potentially all). Then, top genes are selected as markers based on some metric (e.g. fraction of variance explained).

